# Network-based identification of genetic factors in Ageing, lifestyle and Type 2 Diabetes that Influence in the progression of Alzheimer’s disease

**DOI:** 10.1101/482844

**Authors:** Utpala Nanda Chowdhury, Shamim Ahmad, M. Babul Islam, Fazlul Huq, Julian M.W. Quinn, Mohammad Ali Moni

**Affiliations:** Dept. of Computer Science and Engineering, University of Rajshahi, Rajshahi, Bangladesh; Dept. of Applied Physics and Electronic Engineering, University of Rajshahi, Rajshahi, Bangladesh; Decipline of Biomedical Science, Faculty of Medicine and Health, The University of Sydney, NSW, Australia; Garvan Institute of Medical Research, University of New South Wales, Sydney, Australia

## Abstract

**Motivation:** Alzheimer’s disease (AD) is currently incurable and the causative risk factors are still poorly understood, which impedes development of effective prevention and treatment strategies. We propose a network-based quantitative framework to reveal details of the complex interaction between the various genetic contributors to AD susceptibility. We analyzed gene expression microarray data from tissues affected by AD, advanced ageing, high alcohol consumption, type II diabetes, high body fat, high dietary fat, obesity, high dietary red meat intake, sedentary lifestyle, smoking, and control datasets. We developed genetic associations and diseasome networks for these factors and AD using the neighborhood-based benchmarking and multilayer network topology approaches.

**Results:** The study identified 484 genes differentially expressed between AD and controls. Among these, 27 genes showed elevated expression both in individuals in AD and in smoker datasets; similarly 21 were observed in AD and type II diabetes datasets and 12 for AD and sedentary lifestyle datsets. However, AD shared less than ten such elevated expression genes with other factors examined. 3 genes, namely HLA-DRB4, IGH and IGHA2 showed increased expression among the AD, type II diabetes and alcohol consumption datasets; 2 genes, IGHD and IGHG1, were commonly up-regulated among the AD, type II diabetes, alcohol consumption and sedentary lifestyle datasets. Protein-protein interaction networks identified 10 hub genes: CREBBP, PRKCB, ITGB1, GAD1, GNB5, PPP3CA, CABP1, SMARCA4, SNAP25 and GRIA1. Ontological and pathway analyses genes, including Online Mendelian Inheritance in Man (OMIM) and dbGaP databases were used for gold benchmark gene-disease associations to validate the significance of these putative target genes of AD progression.

**Conclusion:** Our network-based methodologies have uncovered molecular pathways that may influence AD development, suggesting novel mechanisms that contribute to AD risk and which may form the basis of new therapeutic and diagnostic approaches.

## 1 Introduction

Alzheimer’s disease (AD) is the most common form of dementia and is characterized by gradual degeneration in memory, cognitive processes, language use, and learning capacity Duthey (2013); Rahman *et al*. (2018b). Initial indications begin with a reduced ability to retain recent memories, but with progression all cognitive functions are inevitably affected, resulting in complete dependency for basic daily activities and greatly increased risk of premature death Serrano-Pozo *et al*. (2011); Rahman *et al*. (2018c). AD is irremediable that accounts for 60% to 80% of all dementia cases and estimated to affect over 24 million people worldwide. In the United States, 93,541 deaths resulting from AD were officially recorded in 2014, which is ranked sixth among all causes of death in the United States and fifth among all causes of death after 65 years of age. The premature death rate of AD sufferers increased by 89% within the five years up to 2010, whereas death rates associated with other major morbidities such as cardiac disease, stroke, breast and prostate cancer, and AIDS all declined in that frame. Currently, one new case of the AD is developed in every 66 seconds, a rapid rate of development expected to double by 2050 Association *et al*. (2017); Rahman *et al*. (2018a).

The pathogenesis of the AD is not clearly understood, but it is hypothesized that both genetic and environmental factors are the primary causes. Genes encoding amyloid precursor protein (APP), presenilin 1 (PSEN1) and presenilin 2 (PSEN2) have been identified as associated with AD development Waring and Rosenberg (2008). Age is found to be the most influential risk factor for AD, along with a sedentary lifestyle. Typically AD develops after the age of 65 years and almost half individuals over 85 years old have AD Lindsay *et al*. (2002). Obesity also increases the risk of dementia and eventually AD Kivipelto *et al*. (2005). Type II diabetes, hypertension, smoking and dietary fats can increase the risk of developing AD Mayeux and Stern (2012) Janson *et al*. (2004) Morris *et al*. (2003); Rahman *et al*. (2018a). Meta-analysis of prospective studies suggests that alcohol consumption in late life yields reduced the risk of dementia and hence reduced the risk of AD Anstey *et al*. (2009). AD is a complex polygenic disorder, and many of the associated factors are yet to be identified. For these reasons, there are many problems with accurate diagnosis, characterizing heterogeneous groups of patients who may respond differently to treatment and complicate decisions regarding effective treatment. With such poor understanding of a disease the discovery of further genetic factors could be an important avenue to development of improved diagnostic profiles, and a clearer understanding of the disease process Tilley *et al*. (1998). The key genetic factors associated with susceptibility to complex diseases can be effectively unravelled by genome-wide association studies, and the usefulness of this approach has been proven empirically. Our methodology employed here aims to identify genetic factors influencing common and complex conditions against the background of the random variation seen in a population as a whole Altshuler *et al*. (2008); Moni and Liò (2015).

Molecular association analyses, including differential gene expression determination, protein-protein interactions (PPIs), gene ontologies (GO) and metabolic pathways can ascribe gene activity-based relationships between AD and various risk factors for the disease Rzhetsky *et al*. (2007) Moni and Liò (2014a). Differentially expressed transcripts seen in studies comparing control individuals with individual affected by a disease (or disease risk factor) identify putative disease-associated genes of interest; a differentially expressed gene identified in AD can be more strongly linked to AD when it is shared with AD risk factor differentially expressed genes Goh *et al*. (2007) Feldman *et al*. (2008); Moni *et al*. (2018). From a proteomics point of view, genes are also associated through biological modules such as PPIs, gene ontologies or molecular pathways Lage *et al*. (2007) Suthram *et al*. (2010); Satu *et al*. (2018).

Recently there has been many advances in network-based integrative analytical methods used by researchers to identify possible roles of biomolecules in complex diseases Moni and Liò (2014b) Moni and Lio’ (2017) Torkamani *et al*. (2008). A number of transcriptomic and genetic studies have been conducted on AD Bertram *et al*. (2010) Logue *et al*. (2011) Seshadri *et al*. (2010); Hossain *et al*. (2018a). However, most of these findings have been limited at the transcript level, since the functional interactions among the gene products have not commonly been considered. Since biological molecules interact with each other to carry out functions in biological processes in cells and tissues, integrative analysis within network medicine context is essential to understand the molecular mechanisms behind diseases and to identify critical biomolecules. Thus, we have used a network-based analysis to determine the genetic influence of associated risk factors and disorders for AD progression, including studies of gene expression profiling, PPI sub-network, gene ontologies and molecular pathways. An extensive study regarding phylogenetic and pathway analysis was therefore conducted to reveal the genetic associations of the AD. Significance of these genes and pathways in AD processes were also validated with gold benchmarking datasets including Online Mendelian Inheritance in Man (OMIM) and dbGaP gene-disease associations databases.

## 2 Materials and Methods

### 2.1 Data

We have analyzed gene expression microarray datasets to identify the association of different factors with the AD at the molecular level. All the datasets used in this study were collected from the National Center for Biotechnology Information (NCBI) Gene Expression Omnibus database (https://www.ncbi.nlm.nih.gov/geo/). Ten different datasets with accession numbers: GSE1297, GSE23343, GSE15524, GSE25941, GSE1786, GSE68231, GSE6573, GSE25220, GSE52553, and GSE4806 were analyzed for this study Blalock *et al*. (2004) MacLaren *et al*. (2010) Raue *et al*. (2012) Radom-Aizik *et al*. (2005) Kakehi *et al*. (2015) Misu *et al*. (2010) Herse *et al*. (2007) Hebels *et al*. (2011) McClintick *et al*. (2014) Büttner *et al*. (2007); Hossain *et al*. (2018b). The AD dataset (GSE1297) is obtained by gene expression profiling of hippocampal tissues on 31 separate microarrays from nine control subjects and 22 AD patients with varying severity. The type II diabetes dataset (GSE23343) contains gene expression data obtained through extensive analysis after conducting liver biopsies in humans. The source of the obesity dataset (GSE15524) is subcutaneous and omental adipose tissue analyzed through expression profiling of 20,000 probes in 28 tissue samples. The advanced age dataset (GSE25941) consists of a global microarray data from skeletal muscle transcriptome of 28 different subjects. The sedentary lifestyle dataset (GSE1786) was obtained by expression profiling array from the vastus lateralis muscle using needle biopsies. The high-fat diet (HFD) dataset (GSE68231) is the expression data from human skeletal muscle identifying accumulation of intramyocellular lipid (IMCL). The high body fat (HBF) dataset (GSE6573) is an Affymetrix human gene expression array data from the abdominal fat tissue. The red meat dietary intervention dataset (GSE25220) is an Agilent-014850 whole human genome microarray data from human colon biopsies before and after participating in a high red-meat dietary intervention. The alcohol consumption dataset (GSE52553) is an Affymetrix human gene expression array data of Lymphoblastoid cells from 21 alcoholics and 21 control subjects. The smoking dataset (GSE4806) is a gene expression profiles of T-lymphocytes from smokers and non-smokers.

### 2.2 Method

Analyzing oligonucleotide microarray data for gene expression is known to be an effective and responsive approach to identify new molecular determinants of human diseases. In this study, we used this methodology along with global transcriptome analysis to investigate the gene expression profiles of the AD with 8 risk factors and type II diabetes. To mitigate the problems involving messenger RNA (mRNA) data comparison using different platforms and experimental set-ups, we normalized each gene expression data for each disease using the Z-score (or zero mean) transformation for both disease and control state Sakib *et al*. (2018). Each sample of gene expression matrix was normalized using mean and standard deviation. The expression value of gene *i* in sample *j* represented by *g_ij_* was transformed into *Z_ij_* by computing

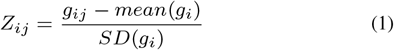

where SD is the standard deviation. Comparing values of gene expression for various samples and diseases are made possible by this transformation. Data were transformed using *log*_2_ and differentially expressed genes for both disease and control states were obtained by performing unpaired student t-test, and significant genes were identified by using threshold values. A threshold for p-value and absolute base two log fold change (logFC) values were set to at most 0.05 and at least 1.0 respectively. We built two infectome-diseasome relationships networks using Cytoscape (v3.5.1) Smoot *et al*. (2010); Moni *et al*. (2014) for both up-regulated and down-regulated genes focusing on the AD. Each node of the networks are either diseases or associative factors. These networks can also be considered as bipartite graphs where diseases or factors are connected when they share at least 1 differentially expressed gene.

We used the web-based visualization software STRING Szklarczyk *et al*. (2016) for the construction and analysis of the Protein-Protein Interaction (PPI) network which was further analyzed by Cytoscape. An undirected graph representation was used for the PPI network, where the nodes indicate proteins and the edges symbolize the interactions between the proteins. We performed a topological analysis using Cyto-Hubba plugin Chen *et al*. (2009) to identify highly connected proteins (i.e., hub proteins) in the network and the degree metrics were employed Calimlioglu *et al*. (2015); Xu *et al*. (2015). For further introspection into the metabolic pathways of the AD, we incorporated the pathway and gene ontology analysis on all the differentially expressed genes that were common between the AD and the other risk factors datasets using the web-based gene set enrichment analysis tool EnrichR Kuleshov *et al*. (2016). In this analysis, the Gene Ontology (GO) Biological Process (BP) and KEGG pathway databases were selected as annotation sources. For statistical significance, the highest adjusted p-value was considered 0.05 to obtain enrichment results. Obtained GO and pathway were further analyzed by Cytoscape. Moreover, two gold bench mark validated datasets, OMIM (https://www.omim.org/) and dbGaP (https://www.ncbi.nlm.nih.gov/gap) were included in our study to validate the principle of our network based approach.

## 3 Results

### 3.1 Gene Expression Analysis

To identify dysregulated genes linked to AD the gene expression patterns from hippocampal CA1 tissues of AD patients were analyzed and compared with normal subject using the NCBI GEO2R online tool (https://www.ncbi.nlm.nih.gov/geo/geo2r/?acc=GSE1297) Blalock *et al*. (2004). 484 genes (p-value at most 0.05 and absolute *log*_2_ fold change value at least 1.0) were found to be differentially expressed compared to healthy subjects where 336 genes were up-regulated and 148 genes were down-regulated.

In order to investigate the relationship of the AD with 8 risk factors and type II diabetes, we performed several steps of statistical analysis for mRNA microarray data regarding each risk factors and disease. Thus, we selected the most significant over and under regulated genes for each risk factor and disease. Our analysis identified a large number of dysregulated genes, namely 958 genes in advanced ageing, 1405 in high alcohol consumption, 824 in high body fat (HBF), 739 in high fat diet (HFD), 381 in obesity, 482 in high dietary red meat, 800 in sedentary lifestyle, 400 in smoking and 1438 in diabetes type II datasets.

The over- and under-expressed genes identified as in common between AD and other risk factors and diseases were also detected through a cross-comparative analysis. The findings demonstrated that AD shares a total of 35, 34, 18, 15, 13, 10, 8, 7 and 4 significant differentially expressed genes with type II diabetes, alcohol consumption, sedentary lifestyle, ageing, HFD, obesity, smoking, HBF and dietary red meat datasets respectively. Two infectome-diseasome associations networks centered on the AD were built using Cytoscape to identify statistically significant associations among these risk factors and diseases. Network shown in Figure-1 interprets the association among up-regulated genes and another network shown in Figure-2 depicts relations between among down regulated genes. Notably, 3 significant genes, HLA-DRB4, IGH and IGHA2 are commonly up-regulated in the AD, type II diabetes and alcohol consumption datasests; 2 significant genes IGHD and IGHG1, were commonly up regulated among the AD, type II diabetes, alcohol consumption and sedentary lifestyle datasets. It is noteworthy that a relatively higher number of differentially expressed genes was identified as in common between the AD and type II diabetes datasets, whereas the AD and high dietary red meat shared only 4 differentially expressed genes.

**Fig. 1.**
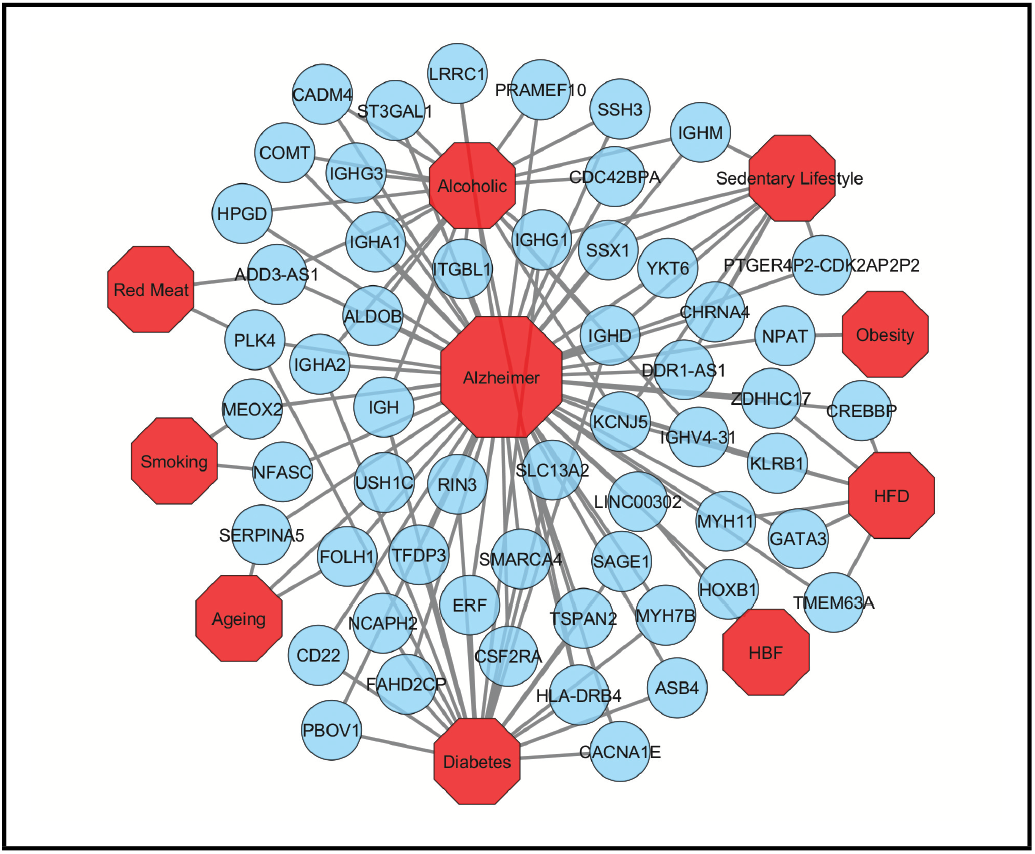
Diseasome network of the AD with type II diabetes, ageing, sedentary lifestyle, HFD, HBF, high dietary red meat, high alcohol consumption, obesity and smoking. Red-colored octagon-shaped nodes represent categories of factors and or disease, and round-shaped sky blue-colored nodes represent up-regulated genes that are common for the AD with the other risk factors and or diseases. A link is placed between a risk factor or disease and gene if alteration expression of that gene is associated with the specific disorder.

**Fig. 2.**
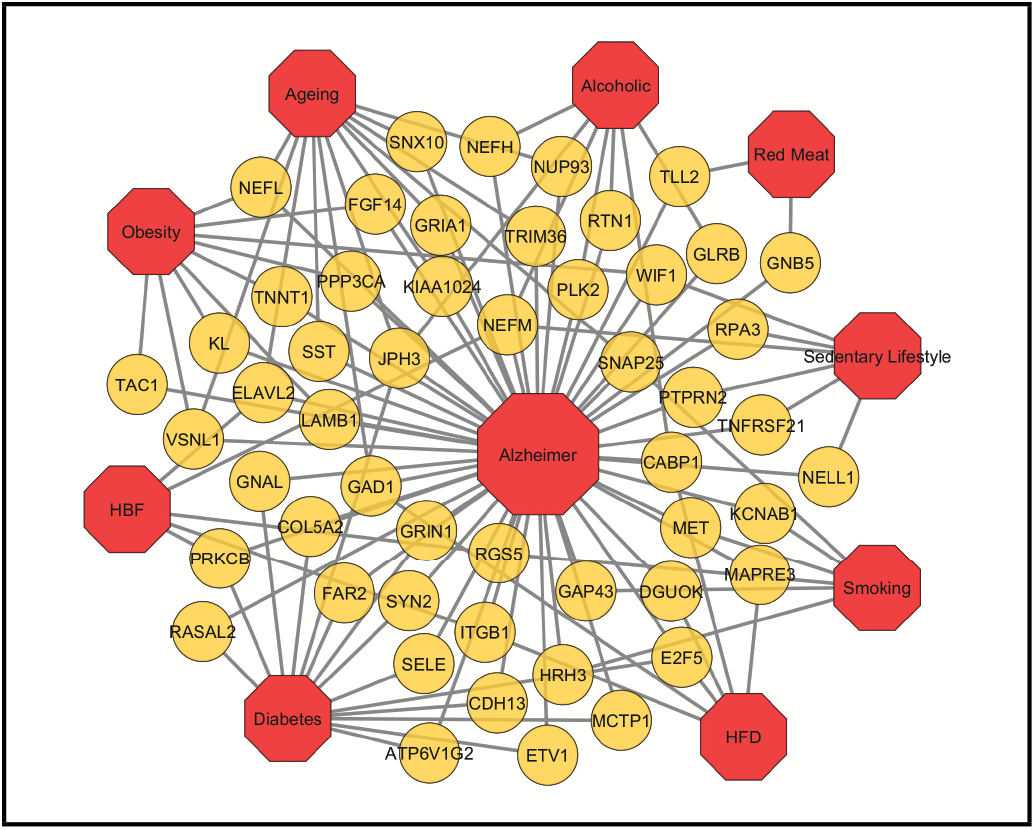
Diseasome network of the AD with type II diabetes, ageing, sedentary lifestyle, HFD, HBF, dietary red meat, high alcohol consumption, obesity and smoking. Red-colored octagon-shaped nodes represent categories of factors and or disease, and round-shaped yellow-colored nodes represent down-regulated genes that are common for the AD with the other risk factors and or diseases. A link is placed between a risk factor or disease and gene if altered expression of that gene is associated with the specific disorder.

### 3.2 Protein-Protein Interaction Network Analysis

The PPI network was constructed using all the distinct 108 (from total 144) differentially expressed genes that were identified as in common between the AD and other risk factors and disease datasets (Figure-3). Each node in the network represents a protein and an edge indicates the interaction between two proteins. The network is also grouped into 9 clusters representing risk factors and diseases to depict the protein links. It is notable that KCNJ5 protein belongs to the highest number (3) of clusters indicating that it is gene most commonly found among the AD, alcohol consumption, HFD and sedentary lifestyle datasets and interacts with other proteins from different clusters. In addition the protein products of PLK4, PRKCB, E2F5, GAD1, VSNL1, RGS5, ITGB1, CABP1 and NEFM belong to two clusters each, and interact with other proteins in the network. For topological analysis, a simplified PPI network was constructed using Cyto-Hubba plugin to show 10 most significant hub proteins (Figure-4), which are CREBBP, PRKCB, ITGB1, GAD1, GNB5, PPP3CA, CABP1, SMARCA4, SNAP25 and GRIA1. The most significantly identified hub protein CREBBP (CREB binding protein) plays major role during the evolution of central nervous system. Alteration of CREBBP activity is known to be implicated in AD progression Rouaux *et al*. (2004).

**Fig. 3.**
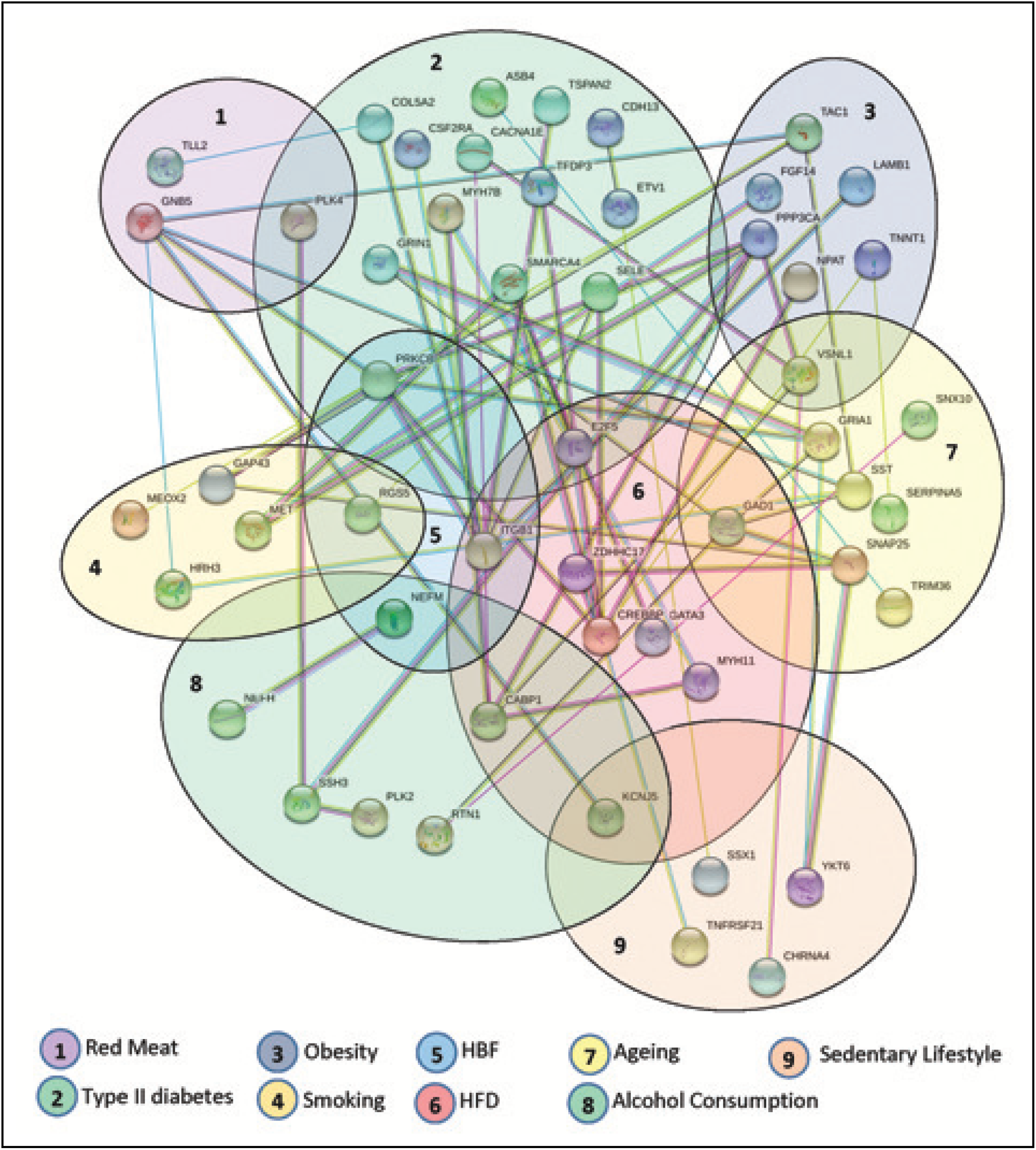
Protein-Protein interaction network of commonly dysregulated genes among AD and other risk factors and diseases. Each cluster indicates the gene belongings.

**Fig. 4.**
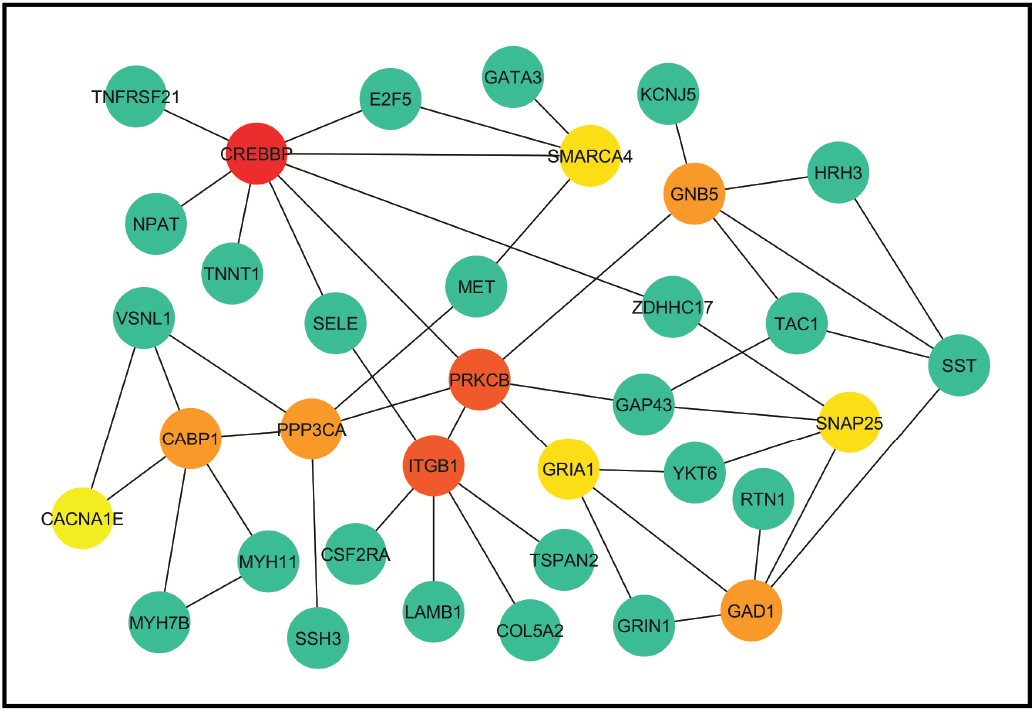
The simplified PPI network of the commonly dysregulated genes among between AD and other risk factors and diseases. The 10 most significant hub proteins are marked as red, orange and yellow respectively.

### 3.3 Pathway and Functional Correlation Analysis

In order to identify the molecular pathways associated with the AD and predicted links to the affected pathways, we performed pathway analysis on all the differentially expressed genes that were common among the AD and other risk factors and diseases using the KEGG pathway database (http://www.genome.jp/kegg/pathway.html) and the web-based gene set enrichment analysis tool EnrichR Kuleshov *et al*. (2016). A total of 115 pathways were found to be over-represented among several groups. Notably, nine significant pathways that are related to the nervous system were found which are Long-term potentiation (hsa04720), Synaptic vesicle cycle (hsa04721), Retrograde endocannabinoid signaling (hsa04723), Glutamatergic synapse (hsa04724), Cholinergic synapse (hsa04725), Serotonergic synapse (hsa04726), GABAergic synapse (hsa04727), Dopaminergic synapse (hsa04728), and Long-term depression (hsa04730). These pathways along with some other common pathways found are shown in Table 1. A gene and pathway association is analyzed by constructing a network for the resulted pathways using Cytoscape (Figure-5).

**Fig. 5.**
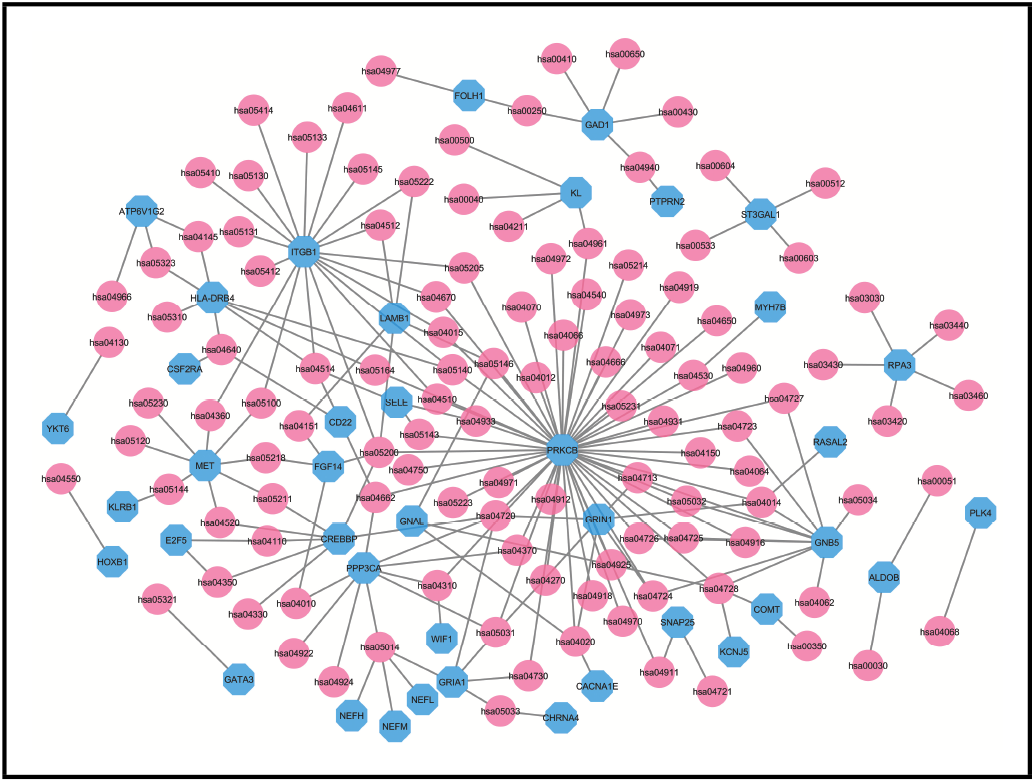
The gene and pathway association network for all pathways obtained for the dysregulated genes common to the AD and other risk factor datasets. Sky blue-colored octagon-shaped nodes represent genes, and round-shaped pink-colored nodes represent pathway (KEGG Id). A link is placed when a gene belongs to a pathway.

**Table 1.** Some significant KEGG pathways that are related to the nervous system and common among the AD and other risk factors and diseases. (Ag=Ageing, T2D=Type II Diabetes, Ob=Obesity, Sm=Smoking, RM=Red Meat, AC=Alcohol Consumption, SL=Sedentary Lifestyle.)

| KEGG ID | Pathway | Genes in pathway | Risk fac./dis. |
| --- | --- | --- | --- |
| hsa04720 | Long-term potentiation | GRIA1, PRKCB, GRIN1, CREBBP, PPP3CA | Ag, Ob, T2D, HBF, HFD |
| hsa05014 | Amyotrophic lateral sclerosis (ALS) | GRIA1, NEFL, NEFM, NEFH, PPP3CA | Ag, AC, HBF, Ob, SL |
| hsa04728 | Dopaminergic synapse | KCNJ5, COMT, GNAL, PRKCB, GNB5 | AC, T2D, HBF, RM |
| hsa05031 | Amphetamine addiction | GRIA1, PRKCB, GRIN1, PPP3CA | Ag, T2D, HBF, Ob |
| hsa04662 | B cell receptor signaling pathway | PRKCB, CD22, PPP3CA | T2D, HBF, Ob |
| hsa04940 | Type I T2D mellitus | GAD1, PTPRN2 | Ag, HFD, SL |
| hsa05100 | Bacterial invasion of epithelial cells | ITGB1, MET | HBF, HFD, Sm |
| hsa05140 | Leishmaniasis | HLA-DRB4, PRKCB, ITGB1 | T2D, HBF, HFD |
| hsa05146 | Amoebiasis | GNAL, PRKCB, LAMB1 | T2D, HBF, Ob |
| hsa00250 | Alanine, aspartate and glutamate metabolism | FOLH1, GAD1 | Ag, HFD |
| hsa04014 | Ras signaling pathway | PRKCB, RASAL2, GRIN1, GNB5 | T2D, RM |
| hsa04310 | Wnt signaling pathway | PRKCB, PPP3CA, WIF1 | HBF, Ob |
| hsa04360 | Axon guidance | ITGB1, MET | HBF, Sm |
| hsa04370 | VEGF signaling pathway | PRKCB, PPP3CA | HBF, Ob |
| hsa04512 | ECM-receptor interaction | ITGB1, LAMB1 | HBF, Ob |
| hsa04514 | Cell adhesion molecules (CAMs) | HLA-DRB4, SELE, CD22, ITGB1 | T2D, HBF |
| hsa04721 | Synaptic vesicle cycle | SNAP25, SNAP25 | Ag, Sm |
| hsa04723 | Retrograde endocannabinoid signaling | PRKCB, GNB5 | HBF, RM |
| hsa04727 | GABAergic synapse | PRKCB, GNB5 | HBF, RM |
| hsa04730 | Long-term depression | GRIA1, PRKCB | Ag, HBF |

We identified over-represented ontological groups by performing gene biological process ontology enrichment analysis using EnrichR on the commonly dysregulated genes between the AD and other risk factors and diseases. Total 215 significant gene ontology groups including peripheral nervous system neuron development (GO:0048935), neurotransmitter transport (GO:0006836), neuromuscular synaptic transmission (GO:0007274), peripheral nervous system development (GO:0007422), negative regulation of neurological system process (GO:0031645), regulation of neurotransmitter secretion (GO:0046928), regulation of neuronal synaptic plasticity (GO:0048168), autonomic nervous system development (GO:0048483), sympathetic nervous system development (GO:0048485), neuromuscular process controlling balance (GO:0050885), neuron apoptotic process (GO:0051402), regulation of neurotransmitter transport (GO:0051588) and neuroepithelial cell differentiation (GO:0060563) were observed (see Table 2). A gene and gene ontology association network is constructed for the obtained gene ontology using Cytoscape (Figure-6).

**Table 2.**
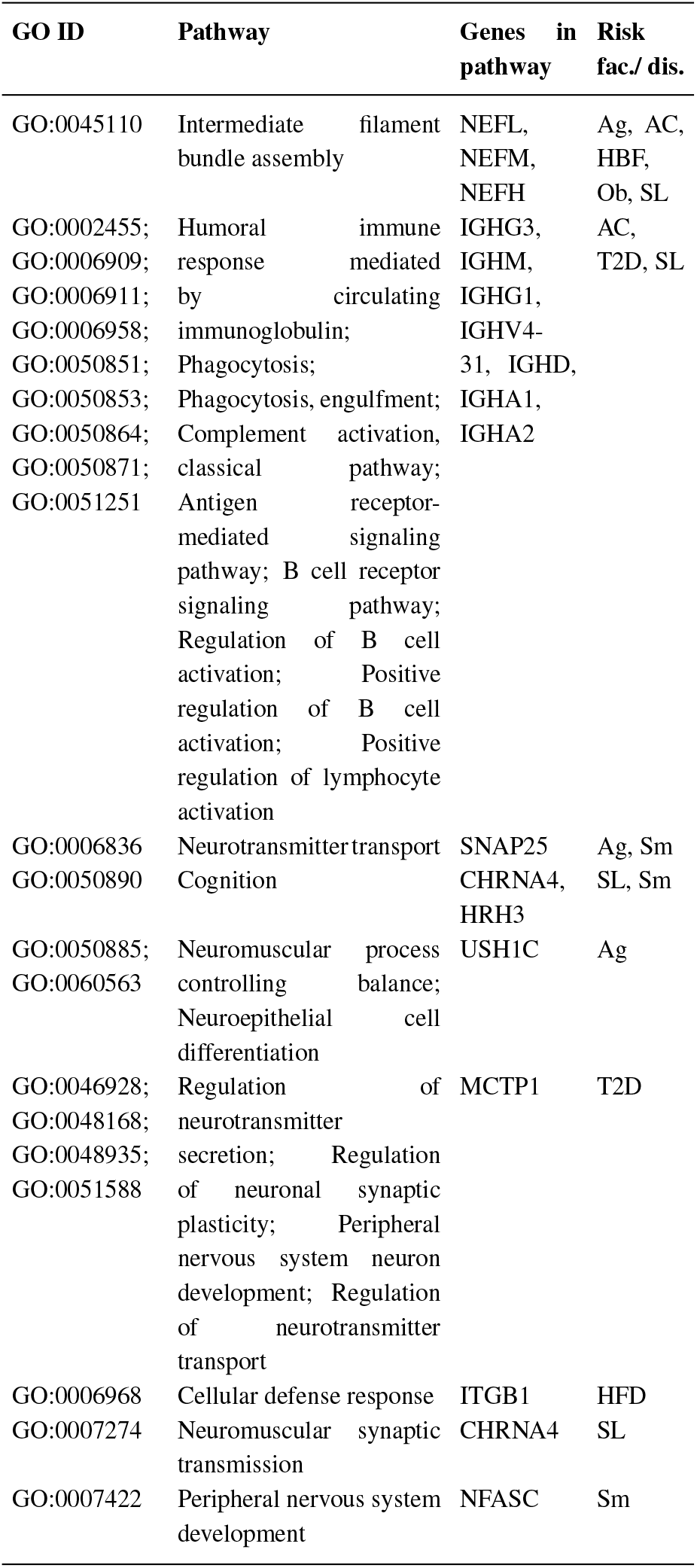
Significant GO ontologies that are related to nervous system, and common between the AD and other risk factors and diseases. (Ag=Ageing, T2D=Type II Diabetes, Ob=Obesity, Sm=Smoking, RM=Red Meat, AC=Alcohol Consumption, SL=Sedentary Lifestyle.)

**Fig. 6.**
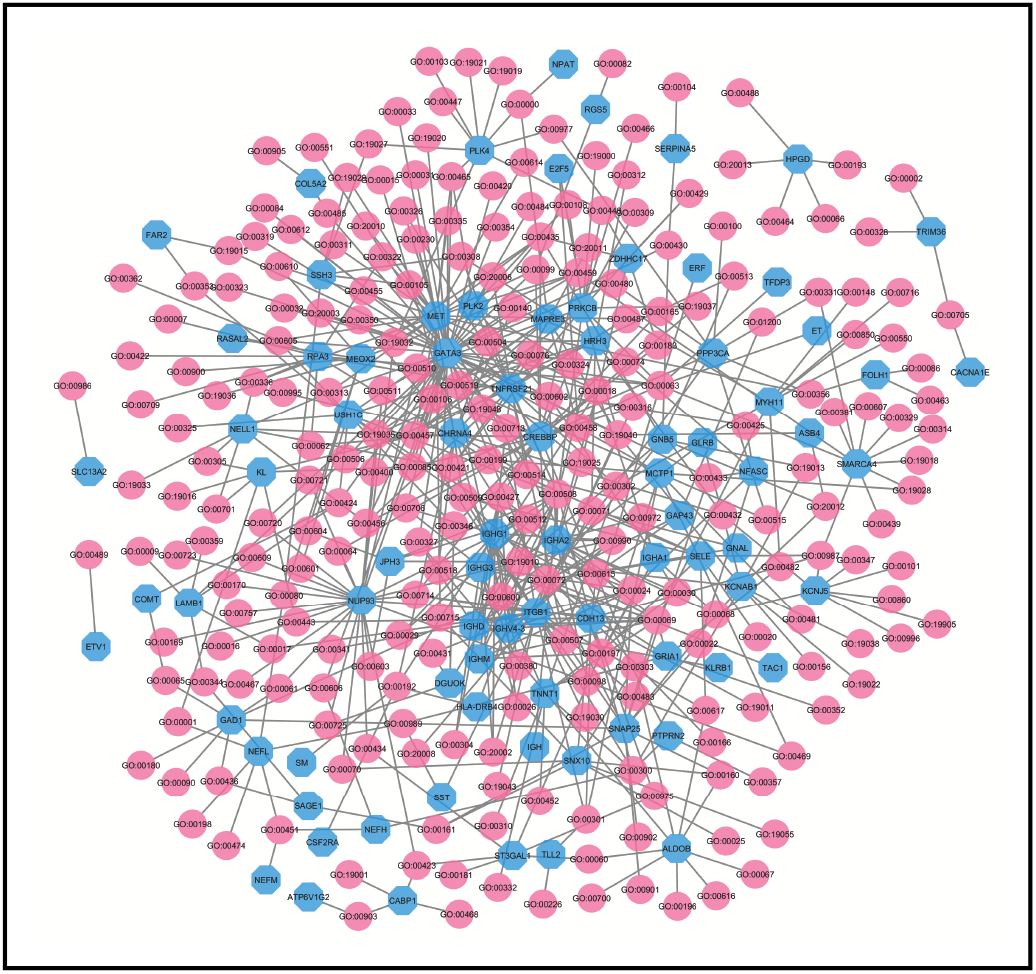
The gene and gene ontology association network for all the gene ontologies obtained for commonly dysregulated genes between the AD and other risk factors and diseases. Sky blue-colored octagon-shaped nodes represent genes, and round-shaped pink-colored nodes represent gene ontology (GO Term). A link is placed when a gene belongs to an ontology.

## 4 Discussion

In this study, we sought novel molecular mechanisms that may affect AD that are made evident by genetic associations with risk factors and diseases that are known to predispose individuals to AD. For this purpose, we conducted analysis in gene expression of AD patients, molecular key pathways, gene ontologies and PPIs. These analyses that employ network-based approach can uncover novel relationships between AD and other susceptibility/risk factor. The findings presented here have not been identified by any previous individual studies. Our study identified several significant genes that may be usefully investigated in other further work, and the hub genes may identify targets for therapeutic interventions in AD. Besides this, our analysis also identified and characterized a number of biological functions related to these genes that throw light on processes that lead to AD.

Our gene expression analysis showed that the AD is strongly associated with type II diabetes (35 genes), alcohol consumption (34 genes), sedentary lifestyle (18 genes) and ageing (15 genes) as they share the maximum dysregulated genes. We constructed and analyzed the PPI network to have a better understanding of the central mechanism behind the AD. For this reason, to construct a PPI network around the differentially expressed genes for our study, we have combined the results of statistical analyses with the protein interactome network. For finding central proteins (i.e., hubs), topological analysis strategies were employed. These identified Hubs proteins might be considered as candidate biomarkers or potential drug targets. From the PPI network analysis, it is observed that 10 hub genes (CREBBP, PRKCB, ITGB1, GAD1, GNB5, PPP3CA, CABP1, SMARCA4, SNAP25 and GRIA1) are involved in the AD.

In addition, disease-related genes play a vital role in the human interactomes via the pathways. In this study, we identified nine significant pathways that are associated with the nervous system which include Long-term potentiation, Synaptic vesicle cycle, Retrograde endocannabinoid signaling, Glutamatergic synapse, Cholinergic synapse, Serotonergic synapse, GABAergic synapse, Dopaminergic synapse, and Long-term depression. Our study also identified several gene ontology groups including peripheral nervous system neuron development, neurotransmitter transport, neuromuscular synaptic transmission, peripheral nervous system development, negative regulation of neurological system process, regulation of neurotransmitter secretion, regulation of neuronal synaptic plasticity, autonomic nervous system development, sympathetic nervous system development, neuromuscular process controlling balance, neuron apoptotic process, regulation of neurotransmitter transport and neuroepithelial cell differentiation. It can be noted that many of these are closely related to the nervous system.

We have also analyzed the differentially expressed genes of each risk factor and type II diabetes with OMIM and dbGaP databases using EnrichR to validate our identified results using the valid gold benchmark gene-disease associations Rana *et al*. (2018). Table 3 shows the genes of each risk factor/disease that are resulted to be associated with the AD. These results corroborate that, the differentially expressed genes of 8 risk factors and type II diabetes are responsible for the AD. As a whole, our findings potentially fill a major gap in understanding AD pathophysiology. They will also open up opportunities to determine the mechanic links between the AD and various risk factors and diseases.

**Table 3.** Gene-disease association analysis of differentially expressed genes of 8 risk factors and type II diabetes with AD using OMIM and dbGaP databases.

| Risk factor/<br>disease | Genes | Adjusted<br>p-value |
| --- | --- | --- |
| Ageing | PLAG1, HMGA2, DNAH11, CR1, VSNL1, DCHS2, F13A1, DISC1, SLC28A1 | 4.34E-02 |
| Alcohol<br>Consumption | GNAQ, GNAS, ARNT, APBB2, ADCY2, ADCY1, IGF1R, MS4A6A, RTN1, ATXN1, PIEZO2, ST3GAL1 | 4.72E-01 |
| Type II<br>Diabetes | AKAP13, HNF4A, HMGA2, BUB1, IGF1R, CR1, DIAPH3, TENM4, CADPS, NEDD9, NPAS3 | 4.66E-01 |
| HBF | PTGIR, THRA, HMGA2, ADCY2, PSEN1, DRD1, BUB1, DBT, PIEZO2 | 1.18E-01 |
| HFD | AKAP13, CREBBP, HNF4A, HMGA2, DNAH11, COL22A1, PIEZO2, RORA, GFRA2, CD33 | 5.17E-01 |
| Obesity | RYR2, RBFOX1, VSNL1, NR2F1, HMGA2 | 1.94E-01 |
| Red Meat | HFE, HMGA2, ADCY2, F13A1 | 4.41E-01 |
| Sedentary<br>Lifestyle | TFAP2A, HSP90AA1, CREB1, THRA, HFE, APOE, TSHR, RYR2, DIAPH3, DCHS2, DBT, CLU | 1.01E-01 |
| Smoking | A2M, OPRD1, SMAD1, ACE, BUB1, CNTNAP2 | 8.45E-02 |

## 5 Conclusions

In this study, transcriptomic data was considered to identify the genetic association of various diseasome relationships with AD. Our findings suggest that these network methods can illustrate disease progression that yields a potential advancement towards having better insight into the origin and development of the AD. Detecting the complex relationship of various risk factors with the disease may disclose novel and useful information for having a better understanding of overall mechanism as well as planning new therapeutic strategies for AD. Using gene expression analysis may be a basis for future accurate disease diagnosis and effective treatment which can be enhanced by the approaches employed in this study. This enhancement may lead to new forms of personalized medicine for increasingly precise insights into disease detection, treatment and remediation for AD.

## Acknowledgements

### Funding

This work has been supported by the… Text Text Text Text.

